# Trajectorygeometry suggests cell fate decisions involve branches rather than bifurcations

**DOI:** 10.1101/2024.02.26.582231

**Authors:** Anna Laddach, Michael Shapiro

## Abstract

Differentiation of multipotential progenitor cells is a key process in the development of any multi-cellular organism and often continues throughout its life. It is often assumed that a bi-potential progenitor develops along a (relatively) straight trajectory until it reaches a decision point where the trajectory bifurcates. At this point one of two directions is chosen, each direction representing the unfolding of a new transcriptomic programme. However, we have lacked quantitative means for testing this model. Accordingly, we have developed the R package TrajectoryGeometry. Applying this to published data we find several examples where, rather than bifurcate, developmental pathways *branch*. That is, the bipotential progenitor develops along a relatively straight trajectory leading to one of its potential fates. A second relatively straight trajectory branches off from this towards the other potential fate. In this sense only cells that branch off to follow the second trajectory make a “decision”. Our methods give precise descriptions of the genes and cellular pathways involved in these trajectories. We speculate that branching may be the more common behaviour and may have advantages from a control-theoretic viewpoint.

## Introduction

Multicellular organisms consist of complex communities of diverse cell types. Remarkably, these all arise from a single cell, the zygote. Accordingly, the development of an organism involves cell fate restriction and differentiation. It has been shown that cell differentiation frequently proceeds via a hierarchy of binary decisions [1], although multifurcations are also possible [2]. Here we will focus on binary cell fate decisions. A typical model of these cell fate decisions is as follows: a multi-potential progenitor cell develops until it reaches a decision point. Here the cell chooses one of two possible fates, each of which requires the initiation of a new developmental programme. After making this choice the cell develops towards the chosen fate. We refer to this as the *bifurcation model* (Figure 1a), where each chosen outcome is seen as more differentiated than the progenitor state.

**Figure 1.**
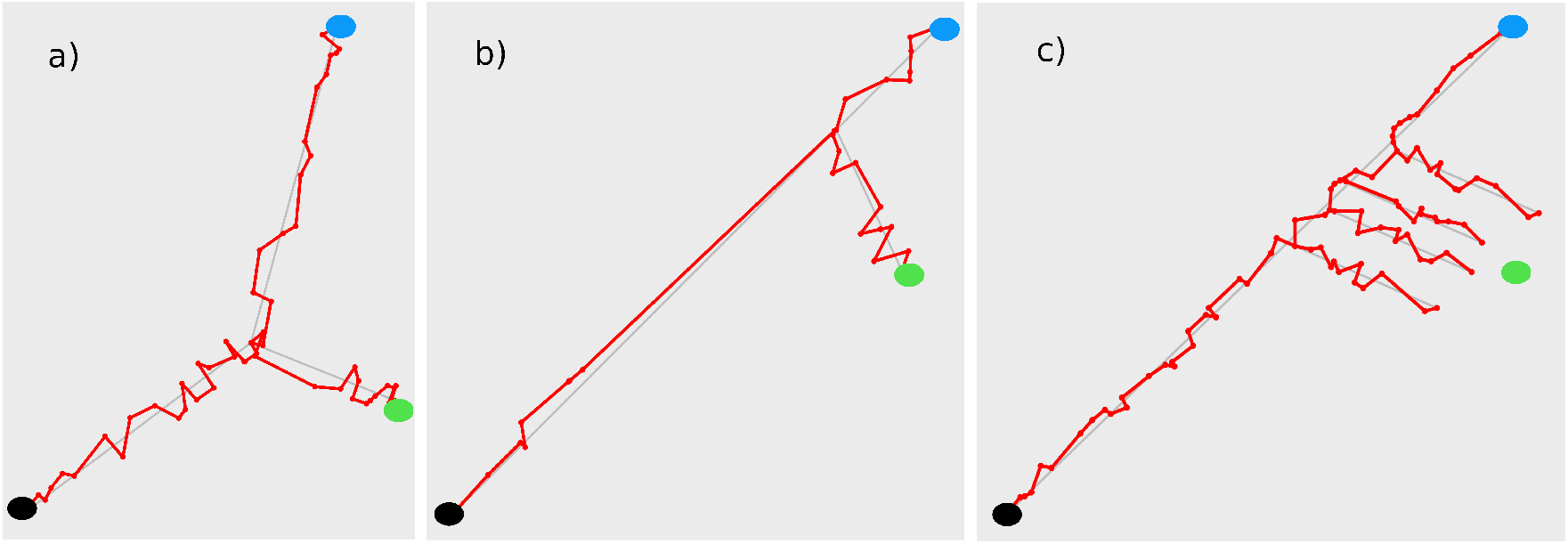
Synthetic data showing bifurcation and branching models. **a)** Bifurcation as a model of cell-fate decision. Bi-potential progenitor cells (black) proceed to a decision point after which they proceed in one of two new directions in gene expression space. **b)** A simplified version of branching behaviour as a model of cell-fate decision. Here bi-potential cells proceed along a default developmental pathway to one of their potential outcomes. The decision is whether or not to leave this default pathway and develop in a new direction. **c)** In this version of branching behaviour, there is a region rather than a single point at which cells choose to leave the default pathway.

Our exploration of lineage decisions in the enteric nervous system (ENS)[3] has uncovered a novel configuration of differentiation trajectories. In contrast to the bifurcation model, enteric gliogenesis forms a default “linear” path of progenitor maturation, from which neurogenic trajectories branch off during embryogenesis. A consequence of this branching configuration (Figure 1b) is that there are no identifiable points of commitment along the gliogenic trajectory and it is only the cells which become neurons that ever “make a decision” and initiate a new developmental program. Further, rather than a single branch point, there seems to be a region along the default trajectory where this branching can take place (Figure 1c). We believe this branching model of lineage decisions allows for plasticity along the default trajectory and underpins the neurogenic potential of mature enteric glial cells.

Having observed this branching model of lineage decisions in the development of enteric neurons and glial cells, we were led to question whether this behaviour is unique to the ENS or whether it might be employed more generally. Here we show that this behaviour is observed in the development of hepatocytes into hepatoblasts and cholangiocytes [4] and the development of postnatal murine olfactory stem cells into into sustentacular cells, neurons and microvillous cells [5].

### TrajectoryGeometry

Our analytic tool for observing default and branching behaviour is our Bioconductor package TrajectoryGeometry[6]. The asynchronicity of most differentiation processes enables the simultaneous profiling of cells at different positions along their developmental trajectory. Whilst many packages exist to infer pseudotime trajectories [7, 8, 9], and the inference of genes that are differentially expressed over pseudotime [10], to our best knowledge TrajectoryGeometry is the first to consider the overall geometry of pseudotime trajectories. The notion of a default trajectory is not new, (see [4] and below) however TrajectoryGeometry provides an analytic footing for this idea by detecting whether a developmental trajectory proceeds in a well-defined direction.

A direction in two dimensions is a point on a circle, e.g., a compass point whereas a direction in three dimensions is a point on the sphere. More generally, a direction in *N* dimensions is a point on the *N −* 1 dimensional sphere, 𝕊^*N−*1^. A path with a well-defined directionality will give rise to points on the circle, the sphere, or 𝕊^*N−*1^ which are tightly clustered around their common center. (See Figure 2.) This allows TrajectoryGeometry to detect whether a differentiation trajectory has a well-defined directionality, to produce a p-value for that directionality and to specify its direction. This direction allows for the identification of genes that are up-and down-regulated along the trajectory and consequently which biological pathways are being up-and down-regulated. Further details are given in [3], in the documentation accompanying [6] and in the supporting information.

**Figure 2.**
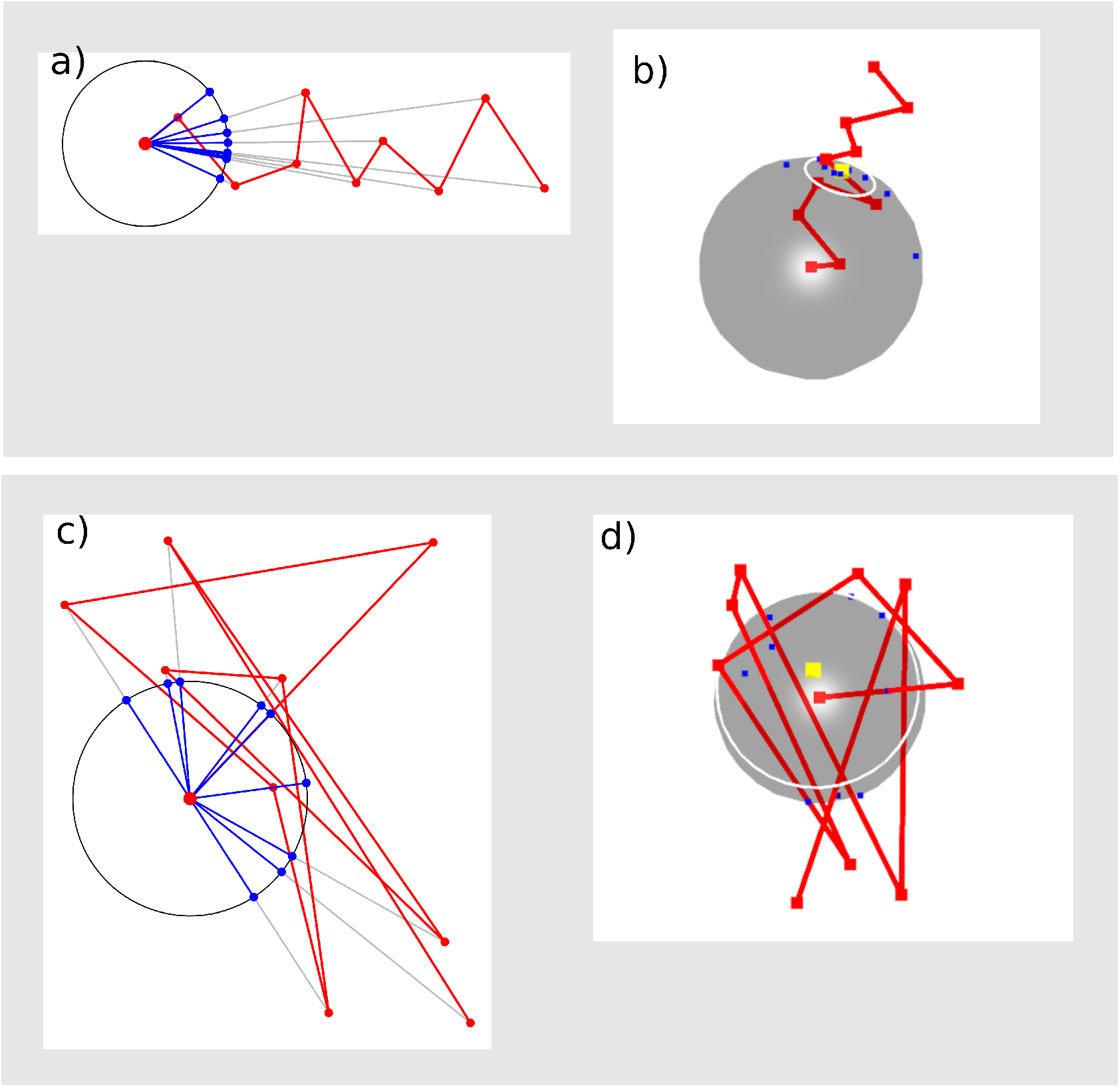
Synthetic data. **a)** A path with a well-defined directionality shown in two dimensions. Directions are sighted from its origin producing a well-clustered set of points on the circle. **b)** The same path shown in three dimensions. Here the directions are the blue points on the sphere. Their common center is shown in yellow. The white circle shows their mean distance from this center. **c)** A path without a well-defined direction. Directions to the points of this path are spread out on the circle. **d)** The same path shown in three dimensions. The white circle showing mean distance to the common center is large, reflecting the lack of common directionality.

## Applications

### Branching cell fate decisions

In the liver, hepatoblasts give rise to hepatocytes and cholangiocytes (Figure 3a). Here we analyse murine single cell data describing this process from [4]. Branching is clearly visible in the 2-dimensional visualisation of pseudotime trajectories, as noted in [4], Figure 3a, where the trajectory that gives rise to cholangiocytes appears to branch off a default time-axis aligned trajectory that ultimately generates hepatocytes. Using TrajectoryGeometry, we see that the small circle observed for the hepatocyte trajectory, in comparison to the larger circle observed for the cholangiocyte trajectory (Figure 3b), suggests hepatocyte development maintains a relatively consistent directionality of gene expression change. Conversely, if the cholangiocyte trajectory is analysed from the decision point onwards, a small circle is also observed, suggesting that it maintains a consistent directionality after branching off (Figure 3b).

**Figure 3.**
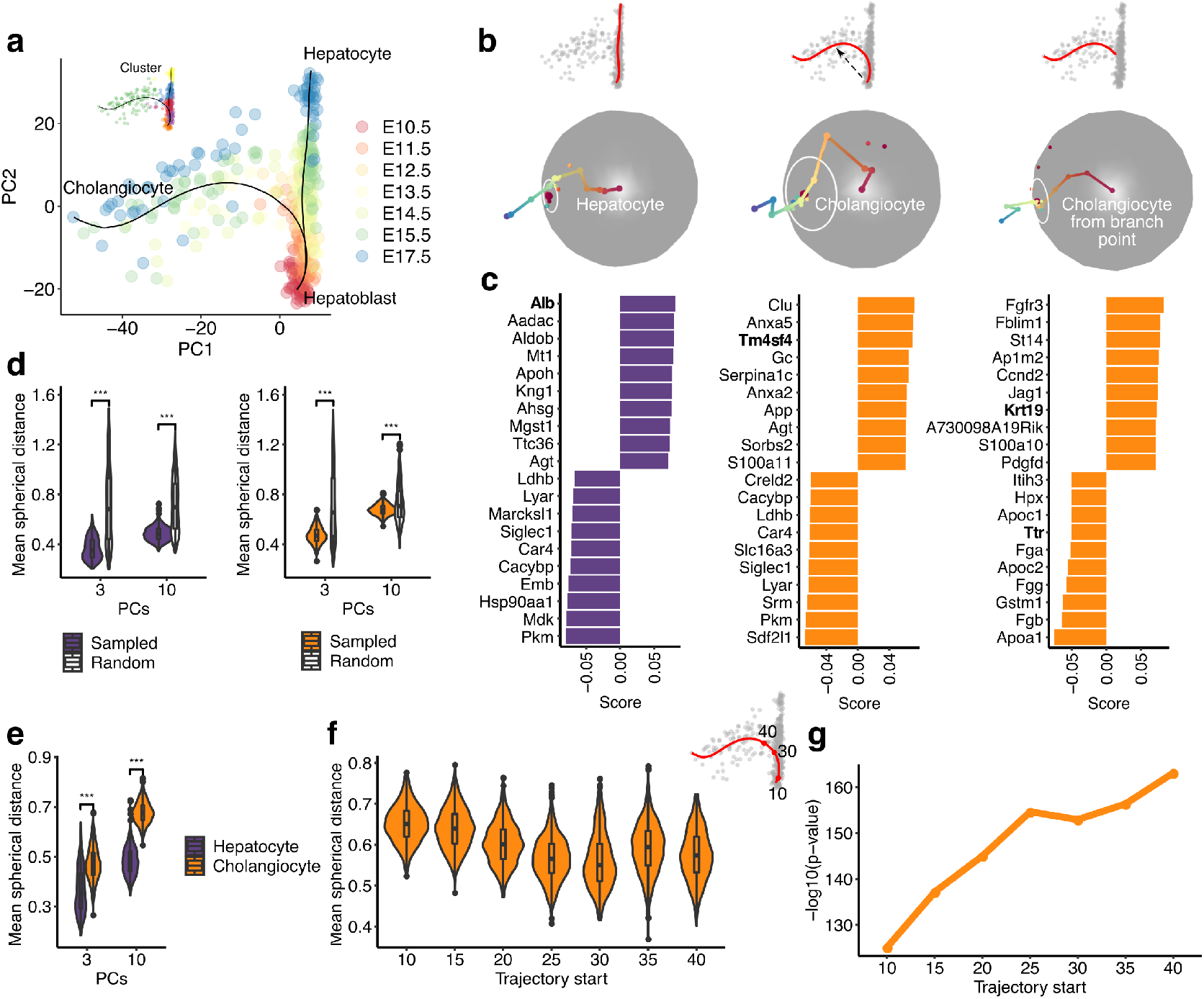
**a)** PCA plot of scRNAseq data for embryonic murine hepatobiliary cells. Pseudotime trajectories inferred using Slingshot are shown on the plot. Cells are coloured by Louvain cluster. **b)** 3-dimensional sampled pathways for hepatoblast to hepatocyte, hepatoblast to cholangiocyte, and decision point to cholangiocyte trajectories together with their projections on the 2-sphere. White circles denote mean distance from center (red dot). **c)** Bar plots showing top 10 up-and downregulated genes for each trajectory as in b). **d)** Violin plots indicating the mean spherical distance (radii of the white circles in b) for paths sampled from the hepatocyte and cholangiocyte trajectories (purple and orange, respectively) relative to random trajectories (white). Statistics calculated using 1000 random paths from each trajectory and the first 3 and the first 10 PCs respectively and the first 3 and the first 10 PCs respectively. **e)** Violin plots indicating the mean spherical distance of the hepatocyte (purple) and cholangiocyte (orange) trajectories. **f)** Violin plots indicating the mean spherical distance for the cholangiocyte trajectory (first 3 PCs) starting from successively later points in pseudotime, as the decision point is approached (30 value on the cholangiocyte trajectory shown in the top right inset). **g)** Line graph indicating the –log10(p-value) for the significance of directionality for the cholangiocyte trajectory (first 3 PCs) relative to random trajectories, starting from successively later points in pseudotime.

Sampling 1000 paths from each trajectory reveals significant directionality in comparison to randomised trajectories for both the cholangiocyte and hepatocyte trajectories (Figure 3d). This is observed whether one uses the first 3 PCs or the first 10 PCs. However direct comparison of cholangiocyte and hepatocyte trajectories reveals that the hepatocyte trajectory maintains a more consistent directionality of gene expression change (Figure 3e)). Furthermore, if the cholangiocyte trajectory segments are analysed starting from successively later points in pseudotime, the mean spherical distance decreases as the decision point is approached and the directionality of the analysed segments becomes more significant (Figure 3f, g), supporting branching behaviour.

Genes positively associated with the directionality of the hepatocyte trajectory include mature hepatocyte markers (e.g. Alb [11]) whereas those negatively associated include markers of hepatoblasts (e.g. Mdk [12]) (Figure 3c). Genes associated with the overall directionality of the cholangiocyte trajectory include those with expression in immature cholangiocytes (e.g. Tm4sf4 [13, 14]) (Figure 3c). This appears to result from the overall directionality being a combination of distinct directionalities before and after the decision point (DP). It is only when we look at the trajectory from the DP to the cholangiocytes that we see markers of mature cholangiocytes (e.g. Krt19 [4]) indicating that a directionality that leads to a cholangiocyte phenotype is not achieved until after branching off (Figure 3c). Mature hepatocyte markers (e.g. Ttr [15]) are amongst genes negatively associated with the DP-cholangiocyte trajectory segment (Figure 3c). This indicates that progenitors have already progressed towards a hepatocyte phenotype when they reach the branch point, and these genes must be subsequently downregulated to acquire a cholangiocyte fate.

Together these findings further support a model where the cholangiocyte trajectory branches off from a default hepatocyte trajectory. Although the inferred trajectory shows a single branch point, the dispersion of cells around this branch, and the fact that the cells are from different embryonic time points, from E11.5 to E14.5, suggest that branching is possible from a continuous section of the “default” trajectory. The change in the directionality of gene expression at the decision point for the cholangiocyte trajectory signifies initiation of a new transcriptomic programme, suggesting that cells are responding to an extrinsic signal. In agreement with this, the cholangiocyte fate decision has been shown to be coordinately regulated by TGF-beta, WNT, Notch and FGF signalling [16, 17, 18, 19, 20, 21] likely in response to factors produced by the periportal mesenchyme.

The liver has remarkable regenerative power, with both cholangiocytes and hepatocytes acting as facultative stem cells able to transdifferentiate if regenerative capacity of the other population is impaired [22]. However, it is interesting to note that hepatocytes, that result from the default trajectory, appear to have unlimited regenerative capacity [23]. It is also interesting that the most abundant cell type (70 % of liver cells are hepatocytes [24]) appears to be produced by default.

### Nested cell fate decisions

Data from [5] offer an opportunity to investigate nested cell fate decisions. Figure 4a), a plot in PCA space, shows postnatal murine olfactory stem cells (also known as horizontal basal cells (HBCs)) giving rise to sustentacular cells, neurons and microvillous cells (MVCs). Visual inspection of this plot suggests that the HBC to Sustentacular trajectory is a default trajectory, consistent with the fact that the latter are produced via direct fate conversion from HBCs. In contrast both neurogenic and MVC trajectories appear to branch off from the sustentacular trajectory at the first decision point (DP1), before diverging from one another at a second decision point (DP2) corresponding to the globose basal cell state.

**Figure 4.**
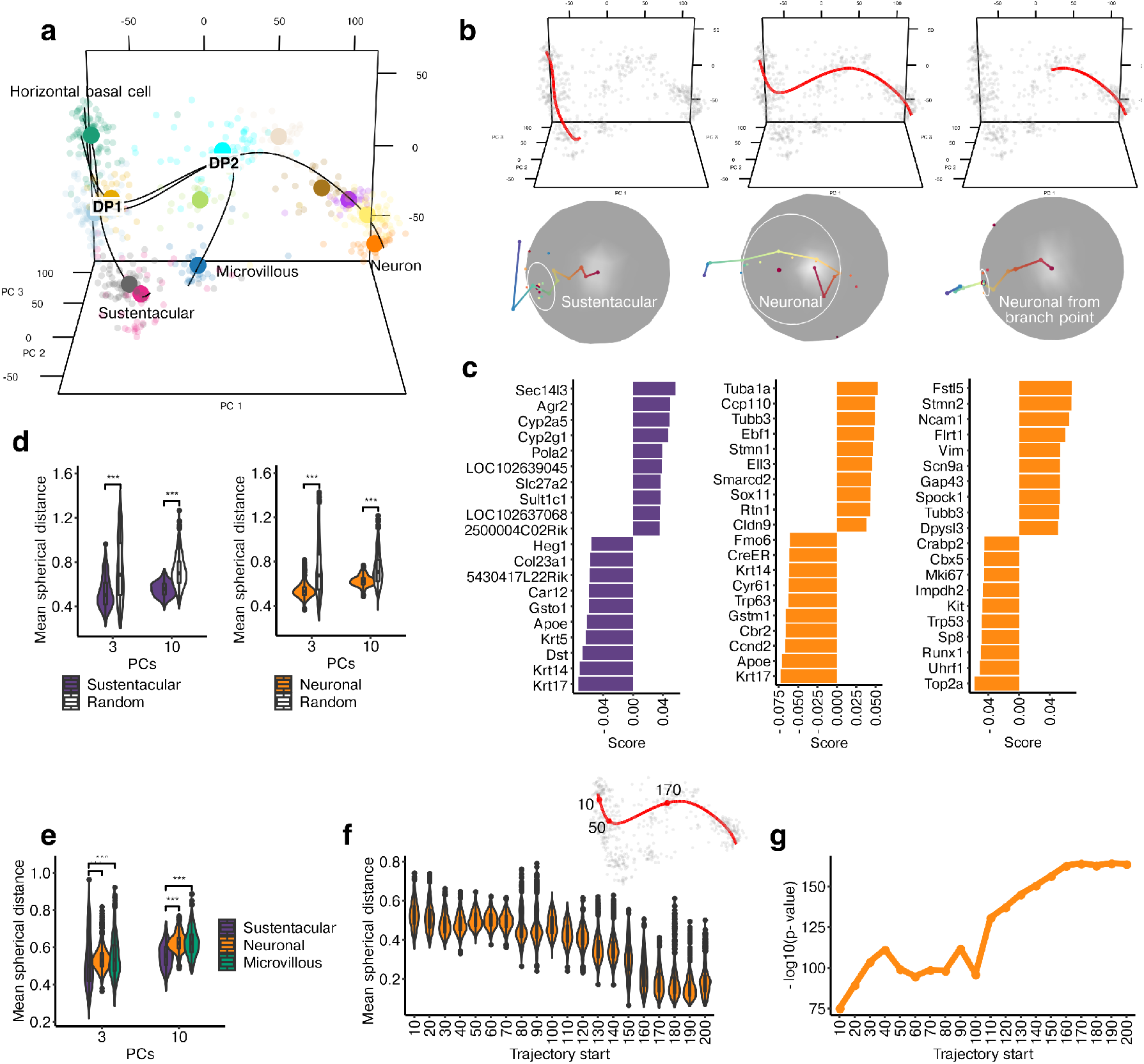
**a)** PCA plot of scRNAseq data for adult murine olfactory cells. Pseudotime trajectories inferred using Slingshot as in [5] are shown on the plot. Cells are coloured by cluster as inferred in [5]. **b)** 3-dimensional sampled pathways for HBC to sustentacular, HBC to neuron, and DP2 to neuron trajectories together with their projections on the 2-sphere. White circles denote mean distance from center (red dot). **c)** Bar plots showing top 10 up-and down-regulated genes for each trajectory as in b). **d)** Violin plots indicating the mean spherical distance (radii of the white circles in b) for paths sampled from the sustentacular and neuronal trajectories (purple and orange, respectively) relative to random trajectories (white). Statistics calculated using 1000 random paths from each trajectory and the first 3 and the first 10 PCs respectively. **e)** Violin plots indicating the mean spherical distance of the sustentacular (purple), neuronal (orange) and microvillous (green) trajectories using 1000 random paths from each trajectory and the first 3 and the first 10 PCs respectively. **f)** Violin plots indicating the mean spherical distance for the neuronal trajectory (first 3 PCs) starting from successively later points in pseudotime, as DP1 and DP2 are approached (values 50 and 170 on the neuronal trajectory shown in the top right inset). **g)** Line graph indicating the –log10(p-value) for the significance of directionality for the neuronal trajectory (first 3 PCs) relative to random trajectories, starting from successively later points in pseudotime.

#### DP1 is a branch point

Focussing initially on the neuronal/sustentacular fate decision, TrajectoryGeometry analysis reveals that although both trajectories show significant directionality in comparison to randomised trajectories (Figure 4d), the sustentacular trajectory displays a more consistent directionality relative to the neuronal (and microvillous) trajectory (Figure 4b,d). Genes with a positive score for the sustentacular trajectory include sustentacular markers (e.g. Cyp2g1 [5]) whereas those with a negative score for the directionality of the sustentacular and neuronal trajectories include HBC stem cell markers (Krt14, Krt5, Trp63 [5] (Figure 4c). Although the top scored genes for the overall neurogenic trajectory HBC-Neurons) include neurogenic markers Sox11 and Tubb3 [25], GO term overrepresentation analysis [26] reveals that the top 5 % of genes with positive scores for this trajectory are highly enriched for cell cycle markers (Figure 5a), suggesting that this directionality does not lead to the mature neuronal phenotype and may be strongly influenced by the proliferative globose basal population at DP2.

**Figure 5.**
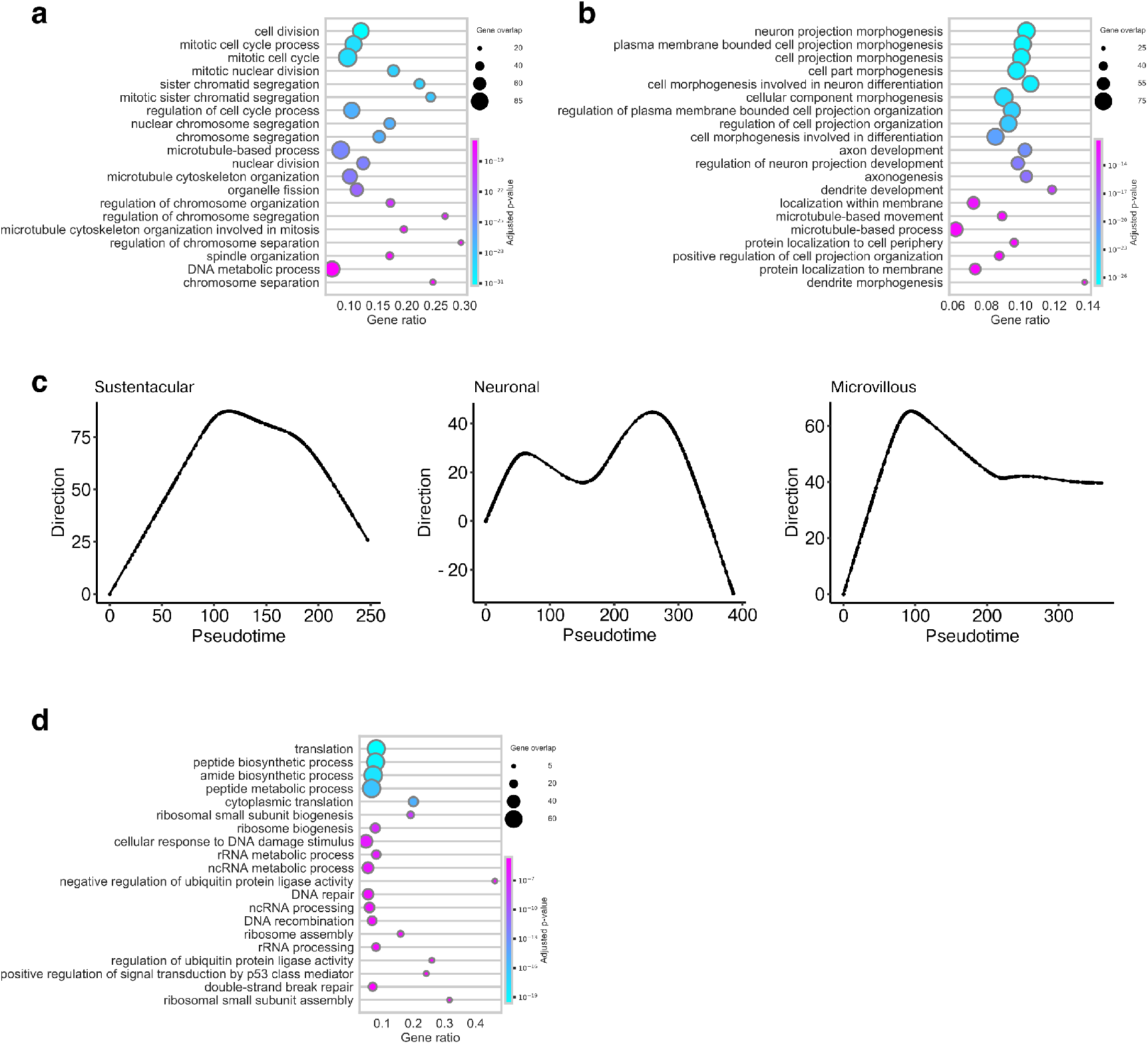
**a)** Dot plot showing GO terms overrepresented among the top 5 % of genes associated with the HBC-neuron directionality. Dot size indicates the overlap for each term, and gene ratio indicates the fraction of genes in each term. **b)** As in a) for genes associated with the DP2-neuron directionality. **c)** Line graphs showing the progress of smoothed trajectories projected onto PC4. **d)** As in a) for the top 5% of genes associated with PC4.

In spite of the relatively more consistent directionality of the sustentacular trajectory, projection onto principle components reveals that it makes a U-turn in PC4 (Figure 5c). Interestingly GO term overrepresentation analysis of the top 5% of genes [26] shows this PC is highly associated with ribosomal genes and protein synthesis (Figure 5d), suggesting transient upregulation of these genes is required for the synthesis of proteins required by the emergent cell type. Indeed, a similar pattern is also seen for the later neuronal (and microvillous) trajectories (Figure 5d), suggesting that this is a common characteristic coincident with differentiation.

Intriguingly, the trajectory from the first decision point to neurons is not straight (see below). Furthermore, Figure 4g shows that the trajectory from progenitors to neurons becomes more directional after passing the first decision point (at value 50 on the neuronal trajectory) and again after passing the second decision point (at value 170 on the neuronal trajectory) (Figure 4b,f,g). Interestingly the top 5 % of genes associated with the segment of the neuronal trajectory from the second decision point onwards (DP2-Neurons) (Figure 4c) are highly enriched [26] in GO terms associated with the development of a mature neuronal phenotype, such as axonogenesis and neuron projection development (Figure 5b). Those negatively associated with the directionality of this post-DP2 trajectory segment include cell cycle markers (e.g. Top2a, Mki67), indicating progress in this direction involves departure from the cycling globose basal cell state at DP2.

#### DP2 exhibits branching behaviour

We now shift our focus to DP2, the neuronal/microvillous decision (Figure 6a).

**Figure 6.**
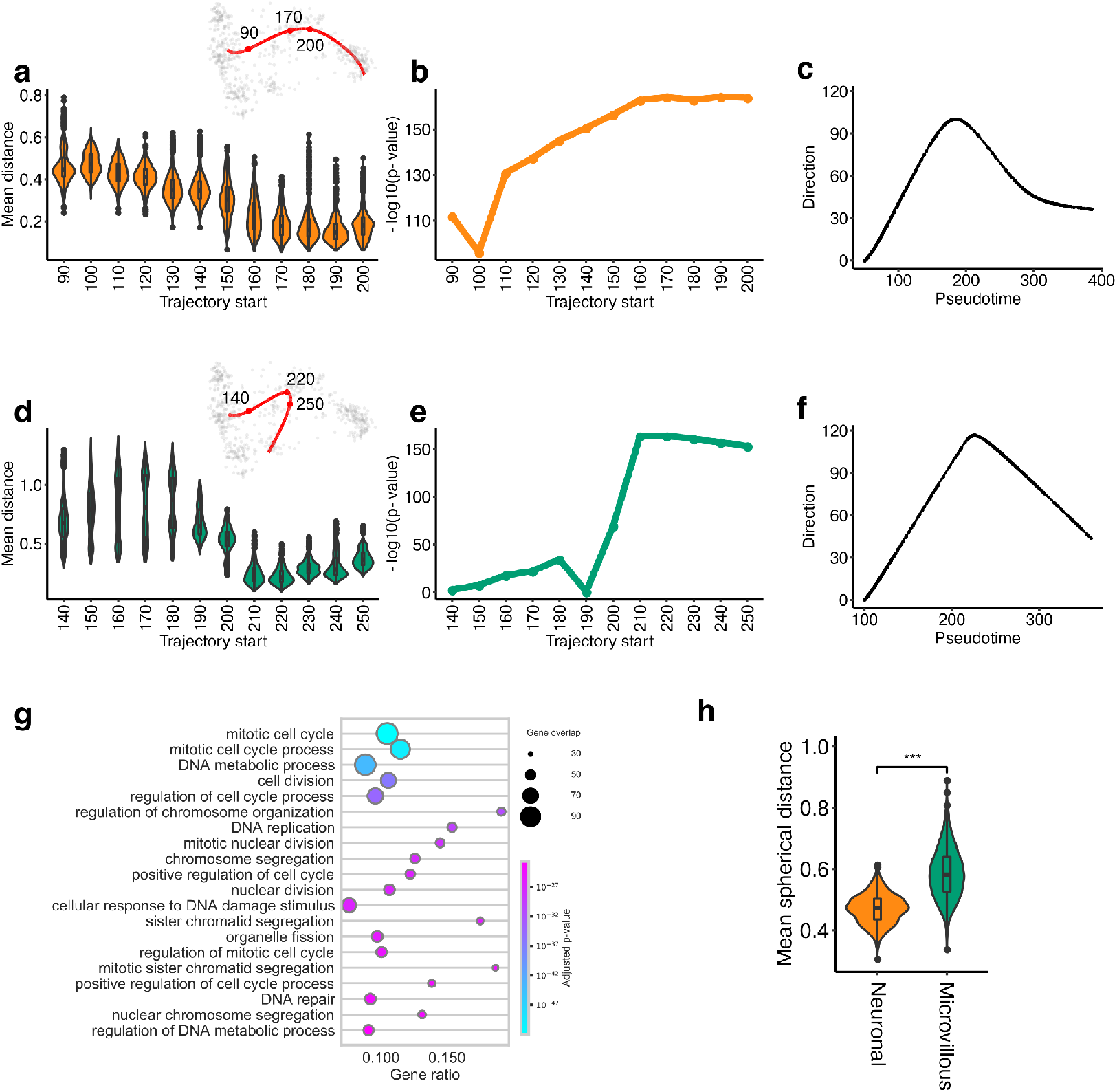
**a)** Violin plots indicating the mean spherical distance for the neuronal trajectory (first 3 PCs) starting from successively later points in pseudotime, as DP2 is approached (value 170 on the neuronal trajectory shown in the top right inset). **b)** Line graph indicating the –log10(p-value) for the significance of directionality for the neuronal trajectory (first 3 PCs) relative to random trajectories, starting from successively later points in pseudotime. **d)** Line graph showing progress of the smoothed neuronal trajectory (available in 5 PCs) projected onto the DP1-DP2 directionality. **d)** Violin plots indicating the mean spherical distance for the microvillous trajectory (first 3 PCs) starting from successively later points in pseudotime, as DP2 is approached (value 220 on the microvillous trajectory shown in the top right inset). **e)** Line graph indicating the –log10(p-value) for the significance of directionality for the microvillous trajectory (first 3 PCs) relative to random trajectories, starting from successively later points in pseudotime. **f)** Line graph showing progress of the smoothed microvillous trajectory (available in 5 PCs) projected onto the DP1-DP2 directionality. **g)** Dot plot showing GO terms overrepresented among the top 5 % of transiently upregulated genes at DP2 for the microvillous trajectory. Dot size indicates the overlap for each term, and gene ratio indicates the fraction of genes in each term. **h)** Violin plots indicating the mean spherical distance of the neuronal (orange) and microvillous (green) trajectories from DP1 onwards using 1000 random paths from each trajectory and PCs 1, 3-10 (omitting PC2).

Specifically we study the microvillous cell (MVC) and neuronal trajectories from DP1 onwards, and subdivide them at DP2 (Figure 4a). Interestingly neither DP1Neurons nor DP1-MVCs is straight and the directionality of both trajectories becomes more significant after DP2 (Figure 6a,b,d,e). Comparing the directions of their initial and final segments (using the first 10 PCs), DP1-Neurons turns approximately 124 degrees and DP1-MVCs makes a turn of 126 degrees (N.B. the first 3 PCs are depicted in Figure 4a). Therefore on initial inspection, DP2 is a bifurcation with each trajectory initiating a new transcriptomic programme.

Notice that by turning more than 90 degrees each of these has partially reversed direction, indicating the partial retraction of a transcriptomic programme. Figure 6c and g show the progression of the pseudotime trajectories for DP1-MVCs and DP1-Neurons in the direction defined by DP1-DP2. Here it can be seen that progress is reversed after the decision point suggesting DP2 is a transient state.

To identify the transiently upregulated genes, we considered the top 5% up-and down-regulated genes in the directions for DP1-DP2, DP2-MVCs and DP2-Neurons and looked at the intersection of the genes that were up-regulated in the first leg with those that were down in the second leg. GO term overrepresentation analysis [26] showed that such genes transiently upregulated for both the microvillous and the neuronal trajectories were highly enriched for cell-cycle associated terms (Figure 6g, Figure 7a), consistent with the proliferative nature of globose basal cells. This suggested that change in direction observed for both neuronal and microvillous trajectories was dominated by reentry into and departure from the cell cycle.

**Figure 7.**
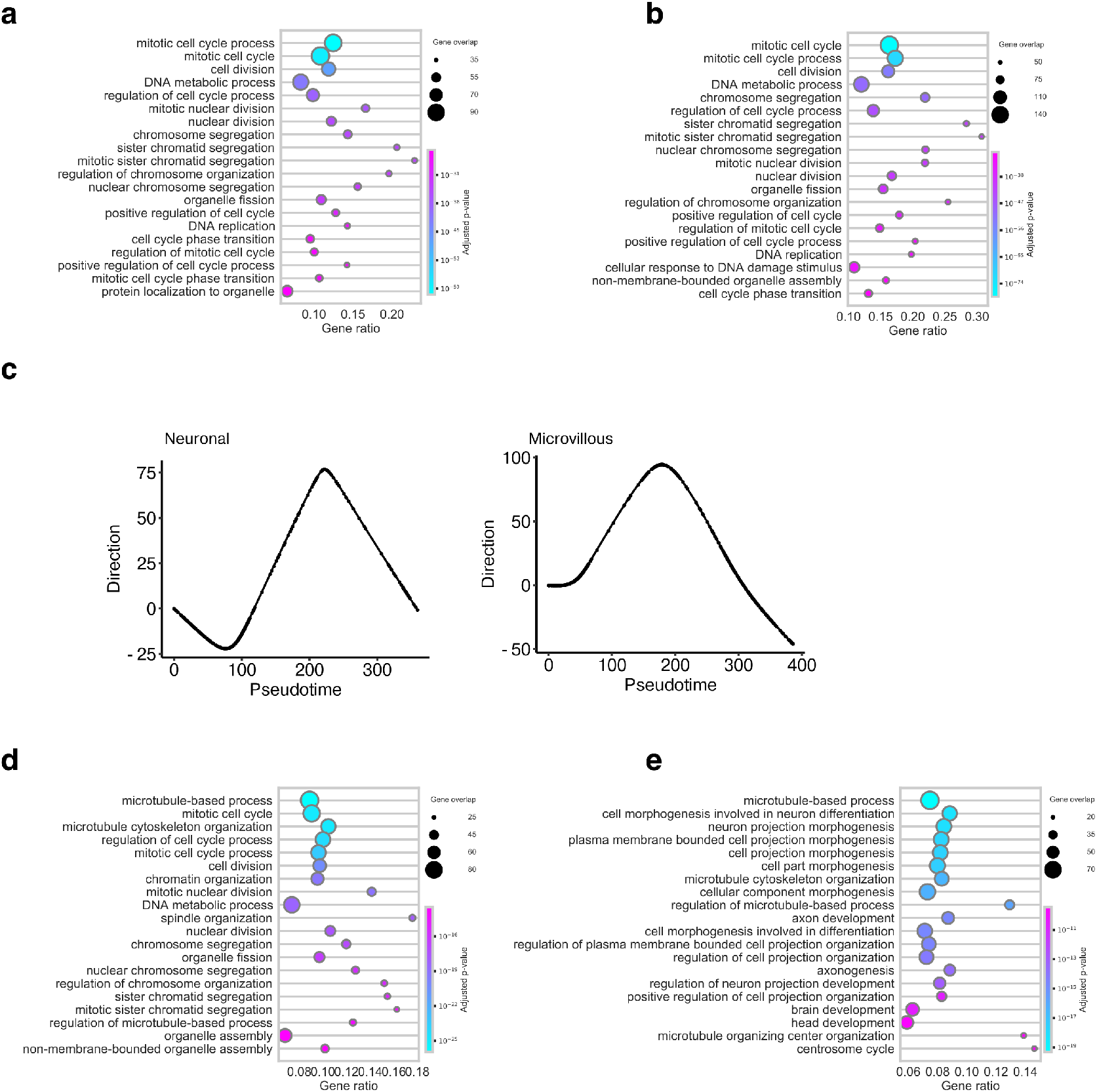
**a)** Dot plot showing GO terms overrepresented among transiently upregulated genes at DP2 for the neuronal trajectory (the intersection of the top 5% up-regulated genes in the DP1-DP2 direction with the top 5% of downregulated genes in the DP2-Neurons direction). Dot size indicates the overlap for each term, and gene ratio indicates the fraction of genes in each term. **b)** As in a) for the top 5% of genes associated with PC2. **c)** Line graphs showing the progress of smoothed trajectories projected onto PC2. **d)** As in a) for the top 5% of genes associated with the DP1-neuron directionality. **d)** As in a) for the top 5% of genes associated with the DP1-neuron directionality omitting PC2.

To test the hypothesis that branching behaviour was being obscured by cellcyle effects, we reanalysed data from DP1 onwards, omitting cell-cycle associated PC2 (Figure 7b, c). Interestingly, this showed the neuronal trajectory to have significantly more consistent directionality than the MVC trajectory (Figure 6h). Therefore if cell-cycle effects are not considered, DP1 appears to be a branch point with the microvillous trajectory branching off from the neuronal trajectory. Put differently, the geometry observed at DP2 results from the transient overlay of cell-cycle on branching behaviour. Furthermore, the top genes associated with the DP1-neuron directionality are enriched in terms that indicate acquisition of a mature neuronal phenotype (e.g. axonogenesis) if PC2 is omitted (Figure 7c, d).

Taken together, these results support the hierarchical branching of trajectories, with the neurogenic and MVC trajectories first branching off a default sustentacular trajectory. Although both MVC and neuronal trajectories then enter the cycling GBC state, the neuronal trajectory appears to be a default upon exit of the cell cycle, whereas the microvillous trajectory branches off, suggesting it may require the input of more extrinsic signals. As microvillous cells are comparatively rare it is parsimonious that these are not produced by default. Importantly, by considering the contribution of individual PCs to directionality we were able to gain insight into the dynamics of cell-type specific transcriptional programmes, and generic transcriptional programmes (translation, cell-cycle.)

## Discussion

In this paper, we have used TrajectoryGeometry to examine the geometry of the cell-fate decisions of multi-potential progenitor cells. Our analyses of several data sets have led us to propose that a branching rather than bifurcating model of cell fate decisions is often employed. In this model, a bipotential progenitor proceeds along a more or less straight *default trajectory* to one of its potential fates. Its other fate arises by *branching off* from this trajectory. In particular, only one of the two cell fates involves initiating a new developmental program. In our experience there is a region along the default trajectory where this branching can take place. Note that development along the default trajectory involves change in gene expression (e.g. the unfolding of a gene expression programme), but not a change in direction of travel in gene expression space as is the case with the initiation of a new programme of gene expression.

Interestingly this model has been anticipated in a more informal manner, e.g. [4] who state: “Thus, the default pathway for hepatoblasts is to differentiate into hepatocytes, but along the way, some hepatoblasts are regulated to differentiate toward the cholangiocyte fate.” TrajectoryGeometry provides the tools to put this in a formal framework by quantifying directionality of trajectories in gene expression space. When it detects directionality, this is expressed as a vector in gene expression space, and this vector tells us which genes are most up-and down-regulated in the developmental process. These allow us to detect the functional pathways involved.

So far we have seen three unequivocal cases of branching behaviour: the branching of neuronal development from the default development of glia[3]; the branching of cholangiocytes from the default development of hepatoblasts into hepatocytes; and the branching of microvillous and neuronal development at DP1 from the default development of horizontal basal cells into sustentacular cells. The point at which microvillous and neuronal development diverges is slightly more complex. Branching behaviour is obscured by a specific process (the cell cycle), which does not exhibit branching behaviour. However, using TrajectoryGeometry we are able to deconvolute process specific effects, and find that this represents another example of branching behaviour.

Note that branching itself is also a transient phenomenon. It seems that there is a region along the default trajectory that is permissive for initiation of the new developmental program. The question arises as to whether branching (or the transient upregulation of cell-cycle) arises due to intrinsic or extrinsic signals. At this point we are unable to reject either of these possibilities. Both cholangiocyte development and neuronal branching at DP1 are known to be responsive to WNT signalling. It is possible that only a portion of the default pathway is responsive to external signals. It is also possible that these external signals only arise at specific developmental time points or in specific cellular environments.

Generalization is premature, but we hypothesize that branching behaviour may be more common than bifurcation and have specific evolutionary advantages. Firstly, we speculate that branching is more robust than bifurcation from a control-theoretic viewpoint. When a cell initiates a new transcriptional program there is always an opportunity for error, both in terms of the signals inducing this change and in terms of the pathways induced by these signals. Changing the transcriptional program of only a subset of the cells exposes fewer cells to this danger. This is particularly advantageous when the branching cell type is required in lower numbers, since in this case only a minority of the cells are required to initiate a new transcriptomic program. We have seen that the minority cell type is the branch outcome in the neurons in the ENS, cholangiocytes in the liver and microvillous cells in the olfactory epithelium. Moreover, we hypothesise that branching behaviour allows for simpler coordinate control of cell numbers for two populations, which is particularly desirable when the two cell types function together (for example glial cells support neurons) and correct proportions must be maintained.

Secondly, branching behaviour may allow for more plasticity in the cells along the default trajectory, as this does not involve initiation of a new transcriptomic programme. This appears to be the case in the ENS where mature glial cells retain neurogenic potential which can be activated under certain conditions. Although both cholangiocytes and hepatocytes retain remarkable plasticity in the liver (both populations are able to transdifferentiate), hepatocytes, generated via the default trajectory, maintain unlimited regenerative capacity.

Thirdly, a default trajectory may allow for the faster generation of a differentiated cell type, particularly in cases where cells undergo direct fate conversion and do not reenter the cell cycle. As an example, sustentacular cells (generated via direct fate conversion) might be urgently required upon loss to maintain the structural integrity of the olfactory epithelium. Furthermore, it has been suggested that sustentacular cells produce crucial factors for olfactory epithelium regeneration [27]; their replenishment might be required before cell types resulting from branching trajectories can be generated.

While the notion of a default trajectory is not new, TrajectoryGeometry gives us the ability to study this analytically, in a way that gives precise measurement of genes and pathways involved in developmental processes. We believe this capability can both provide new answers and raise new questions in the study of developmental biology and cell differentiation.

## Supporting information

We give an overview of the geometry which enables our analysis. This has been previously described in [3] and in the documentation of [6].

### Directionality in ℝ^*n*^

We would like to analyse whether a given developmental trajectory proceeds in a well-defined direction. This breaks down into two questions: first, does it *have* a well-defined direction and second, does it *proceed* in that direction. We have illustrated the first of these questions in Figure 2. In Figure 2 **a)** and **b)**, we see a path with a well-defined directionality, in panels **c)** and **d)** we see a path without a well-defined directionality. These two situations are reflected in the clustering or otherwise of the points on the circle and sphere. More generally, in *N* dimensional space, ℝ^*N*^, a direction is a point on the *N −* 1 dimensional sphere 𝕊^*N −*1^.

Suppose now that we are working with the first 15 principal components (PCs) of gene expression, so that we are working in ℝ^*N*^, *N* = 15. Suppose we now sample 10 cells from a particular trajectory. Call these **c**_1_, …, **c**_10_. Using our first 15 PCs, we then have points **x**_1_, …, **x**_10_ in ℝ^15^. Sighting from the first point **x**_1_ to each of the successive points **x**_2_, …, **x**_10_ gives us 9 directions in R^15^, i.e., 9 points on 𝕊^14^. Call these points **p**_1_, …, **p**_9_. To measure the directionality of the path **x**_1_, …, **x**_10_ (and therefore the directionality of the pseudotime trajectory) we measure how closely **p**_1_, …, **p**_9_ cluster on 𝕊^14^. That is, we find a center *c* on 𝕊^14^ that minimizes mean spherical distance to these points. Charmingly, this measure of directionality can be seen as the radius of an *N −* 2-sphere on 𝕊^*N−*1^.

### Testing for directionality

Let us take *r* to be the minimal mean radius found in the previous paragraph. We can think of *r* as a function of the original trajectory

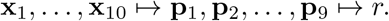

Each time we sample the pseudotime trajectory, we will get a different value for this radius. In this way, repeated sampling (say 1000 times) produces a distribution of estimates of the directionality of the pseudotime trajectory, *r*_1_, …, *r*_1000_. We would now like to use this to estimate a p-value for the directionality of this trajectory.

In order to do this, we turn to permutation testing. That is, we produce (say 1000) randomized trajectories 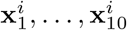 and for each of these 1000 randomized trajectories we compute

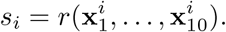

If we detect significant directionality we can proceed to estimate a trajectory’s direction in gene expression space we can use the same method to estimate the direction. Having sampled the trajectory 1000 times, we find 1000 centers, *c*_1_, …, *c*_1000_, each the center for one of the sampled trajectories. We take their common center *ĉ* to give the direction of the overall trajectory. If we are working with the first 10 PCs, this will be a unit vector in ℝ^15^. This can then be translated back into gene expression space where each coordinate represents a gene. The most positive and most negative coordinates in this vector reveal the genes which are most strongly up- and down-regulated along this trajectory.

There are multiple ways of producing a randomized trajectory. The most conservative of these is via matrix permutation. We can consider the path **x**_1_, …, **x**_10_ as a matrix *X* where each of these points is a row of *X*. Thus *X* has dimension 10 *×* 15. We can then independently randomly permute the entries in each column of *X* thus independently permuting the time order for each PC. Further details and other methods of producing randomized paths can be found in [3] and in the vignette accompanying TrajectoryGeometry [6] on Bioconductor.

### Data and code availability

TrajectoryGeometry is available as an R Bioconductor package (10.18129/B9.bioc.TrajectoryGeometry). Data describing the development of hepatocytes into hepatoblasts and cholangiocytes [4] was obtained from GEO under the accession GSE90047 and data describing the development of postnatal murine olfactory stem cells into sustentacular cells, neurons and microvillous cells was obtained from GEO under the accession GSE95601. Scripts to reconstruct olfactory trajectories as presented in [5] were obtained from https://github.com/rufletch/p63-HBC-diff. Scripts used to construct hepatoblast trajectories are available at https://github.com/AnnaLaddach/TrajectoryGeometryData.

